# High-order areas and auditory cortex both represent the high-level event structure of music

**DOI:** 10.1101/2021.01.26.428291

**Authors:** Jamal A. Williams, Elizabeth H. Margulis, Samuel A. Nastase, Janice Chen, Uri Hasson, Kenneth A. Norman, Christopher Baldassano

## Abstract

Recent fMRI studies of event segmentation have found that default mode regions represent high-level event structure during movie watching. In these regions, neural patterns are relatively stable during events and shift at event boundaries. Music, like narratives, contains hierarchical event structure (e.g., sections are composed of phrases). Here, we tested the hypothesis that brain activity patterns in default mode regions reflect the high-level event structure of music. We used fMRI to record brain activity from 25 participants (male and female) as they listened to a continuous playlist of 16 musical excerpts, and additionally collected annotations for these excerpts by asking a separate group of participants to mark when meaningful changes occurred in each one. We then identified temporal boundaries between stable patterns of brain activity using a hidden Markov model and compared the location of the model boundaries to the location of the human annotations. We identified multiple brain regions with significant matches to the observer-identified boundaries, including auditory cortex, medial prefrontal cortex, parietal cortex, and angular gyrus. From these results, we conclude that both higher-order and sensory areas contain information relating to the high-level event structure of music. Moreover, the higher-order areas in this study overlap with areas found in previous studies of event perception in movies and audio narratives, including regions in the default mode network.

**Significance Statement:** Listening to music requires the brain to track dynamics at multiple hierarchical timescales. In our study, we had fMRI participants listen to real-world music (classical and jazz pieces) and then used an unsupervised learning algorithm (a hidden Markov model) to model the high-level event structure of music within participants’ brain data. This approach revealed that default mode brain regions involved in representing the high-level event structure of narratives are also involved in representing the high-level event structure of music. These findings provide converging support for the hypothesis that these regions play a domain-general role in processing events occurring over long timescales.

## Introduction

Recent work has demonstrated that the brain processes information using a hierarchy of temporal receptive windows, such that sensory regions represent relatively short events (e.g., milliseconds to seconds) and higher-order regions represent longer events (e.g., minutes) while inheriting some of the lower-level structure from sensory regions (Baldassano et al., 2017; Chen et al., 2017; Uri Hasson et al., 2015). For example, Baldassano and colleagues (2017) used a hidden Markov model (HMM) to find transitions between stable patterns of neural activity in BOLD data acquired from participants that watched an episode of the TV series Sherlock. The HMM temporally divides data into “events” with stable patterns of activity, punctuated by “event boundaries” where activity patterns rapidly shift to a new stable pattern. They found that, in sensory regions such as early visual cortex, the data were best-fit by a model with short-lasting chunks, presumably corresponding to low-level perceptual changes in the episode; by contrast, when they applied the model to data from a higher-order area such as posterior medial cortex, the best-fitting model segmented the data into longer-lasting chunks corresponding to more semantically meaningful scene changes. Critically, human annotations of important scene changes most closely resembled the model-identified boundary structure found in frontal and posterior medial cortex, which are key hubs in the brain’s default mode network (DMN) (Raichle et al., 2001; Shulman et al., 1997). Studies have also found that the same event-specific neural patterns are activated in default-mode regions by audiovisual movies and by verbal narratives describing these events (Baldassano et al., 2017, 2018; Zadbood et al., 2017), providing further evidence that these regions represent the underlying meanings of the events and not only low-level sensory information.

Jackendoff & Lerdahl (2006) suggest that music and language are structured into meaningful events that help people comprehend moments of tension and relaxation between distant events. If music resembles language in this way, then the representation of hierarchical event structure in music (e.g., at the level of phrases, sections, and entire songs) and in verbal and audiovisual narratives may be supported by similar neural substrates. Indeed, some evidence already exists for shared neural resources for processing music and language (Jantzen et al., 2016; Koelsch, 2011; Koelsch et al., 2002; Lee et al., 2019; Patel, 2011; Tallal & Gaab, 2006; Asano et al., 2021, Fedorenko et al., 2009; Peretz et al., 2015; Tillmann, 2012). This connection between music and language is also supported by recent behavioral studies showing that instrumental music has the capacity to drive shared narrative engagement across people (Margulis et al., 2019, Margulis et al., 2021, McAuley et al., 2021). In the current work, we test the hypothesis that default mode network (DMN) regions, which represent high-level event structure in narratives, also play a critical role in representing high-level event structure in music.

In our paradigm, we presented fMRI participants with examples of complex real-world music belonging to genres familiar to our participant population: jazz and classical. A separate group of behavioral participants were asked to annotate meaningful events within each of the excerpts. Using a whole-brain searchlight method, we applied hidden Markov models to measure event structure represented in cortical response patterns throughout the brain. The goal of this analysis was to identify brain regions that chunk the stimuli in a way that matched the human annotations. By fitting the model at each ROI and then comparing the observed boundary structure to that of the annotators, we show that – in a group of passive listeners – regions in the default mode network and also sensory areas are involved in representing the high-level event structure in music (i.e., these regions show neural pattern shifts that line up with human annotations of event boundaries). We also show that these event representations become coarser as they propagate up the cortical processing hierarchy.

## Materials and Methods

### Participants

We collected fMRI data from a total of 25 participants (12 female, ages 21–33). We also recruited 7 human annotators for a separate behavioral task (described below). Thirteen of the fMRI participants were native English speakers. The experimental protocol was approved by the Institutional Review Board of Princeton University, and all participants gave their written informed consent.

### Stimuli

Sixteen musical excerpts were selected, based on the criterion that changes between sub-sections would likely be recognized by people without formal music training (e.g., change from piano solo to drum solo). Excerpts also had to be instrumental (i.e. lack vocals). Excerpts were drawn from two different genres (8 classical and 8 jazz). Excerpts were then randomly selected to be truncated (with the introductions kept intact) to one of four different durations: 90 s, 135 s, 180 s, and 225 s, such that there were four excerpts of each length. Furthermore, two excerpts of each duration were sampled from each genre. For example, only two classical excerpts had a duration of 90 seconds and only two jazz excerpts had a duration of 90 seconds. The total duration of the playlist was approximately 45 minutes and there were no breaks between excerpts.

## Experimental design and statistical analysis

The experiment took place over three consecutive days (Figure 1): On the first two days participants heard a playlist of sixteen musical excerpts (once for each day) and on the third day they heard the same playlist for two separate runs while we recorded changes in their BOLD activity using fMRI. Altogether, each participant heard the playlist four times. Each time that a given participant heard the playlist, the excerpts were presented in a different order. However, within a given phase of the experiment (e.g., the first scanner run on day 3), the order of excerpts was kept the same across participants. To promote stable representations of the music, participants listened to the playlist on each of the two days prior to scanning. During these listening sessions, we collected ratings from participants about their enjoyment, engagement, and familiarity with each piece (only familiarity ratings are discussed in this manuscript); these ratings were collected immediately after hearing each piece. Answers for each rating category were given on a 5-point Likert scale where a 1 corresponded to very unfamiliar and a 5 corresponded to very familiar. We found an increase in average familiarity from day 1 to day 2 (t(22)= 9.04, p < 0.0001) indicating that participants remembered the music played in the first pre-scan session. Two participants were excluded from this analysis because their day 2 ratings were lost.

**Figure 1.**
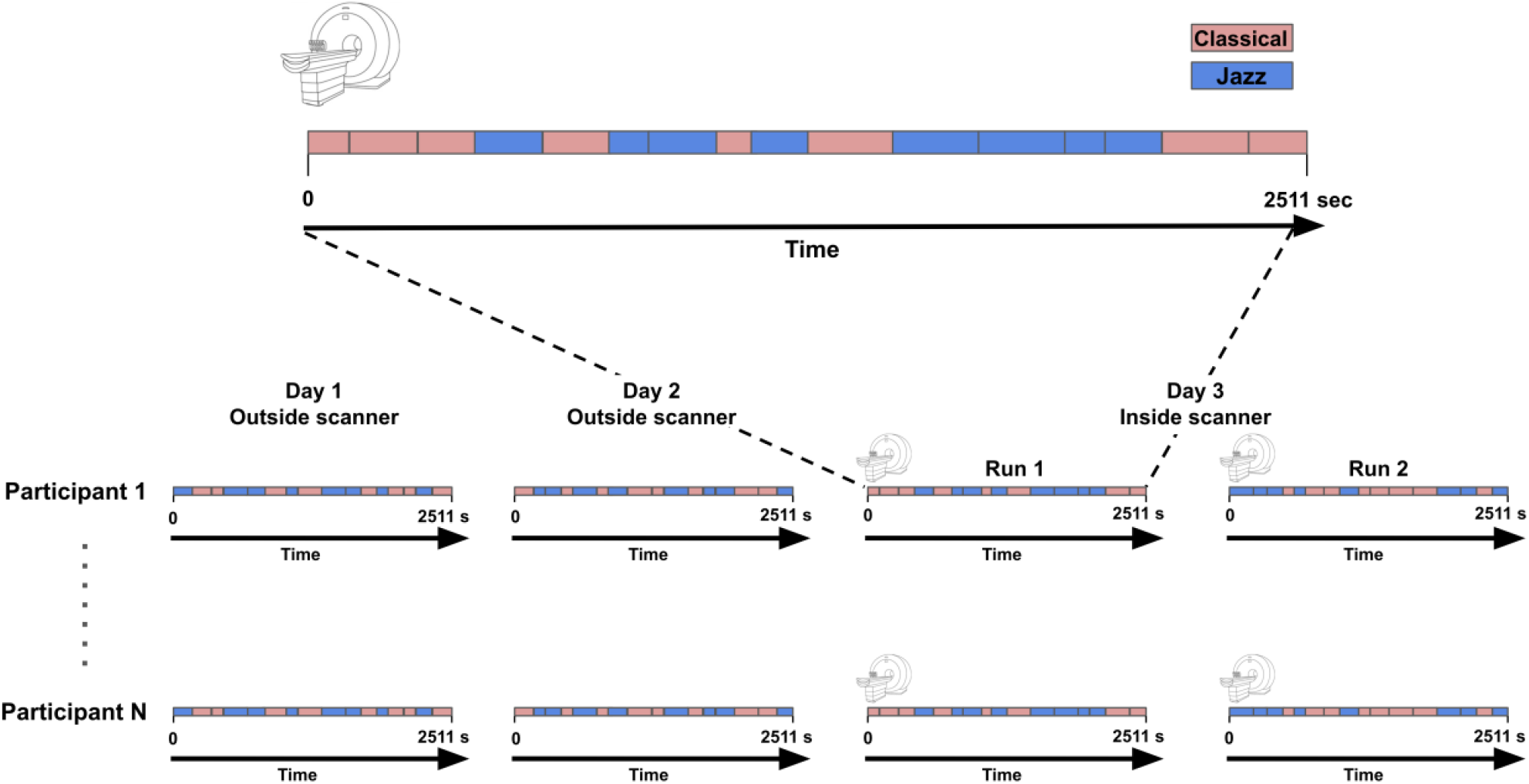
Top: Example 45-minute scanning run with classical excerpts depicted in pink and jazz excerpts in blue. Each block in the timeline represents an excerpt and block lengths reflect excerpt durations. Bottom: Overview of experiment. Participants heard the playlist four times (once on each of the two days prior to scanning and twice on the third day while being scanned). The excerpts were presented in a different order each of the four times that a given participant heard the playlist, but – within a given phase of the experiment (e.g., Run 1 on Day 3) – the order of excerpts was kept the same across participants.

After each of these listening sessions, participants took a short recognition test where they heard 32 randomly drawn 3-second clips of a piece that were either from the actual listening session or a lure (i.e. different piece by the same artist) and made a response using a 5-point Likert scale indicating whether they recognized the excerpt as having been presented previously. In addition to the familiarity ratings across the two pre-scan days, this measure helped us determine if participants had learned the music after each behavioral listening session. Participants showed above-chance discrimination (i.e., higher recognition scores for presented excerpts vs. lures) on both days (Day 1: t(24) = 12.2, p < 0.0001; Day 2: t(24) = 15.1, p < 0.0001; Figure 2).

**Figure 2.**
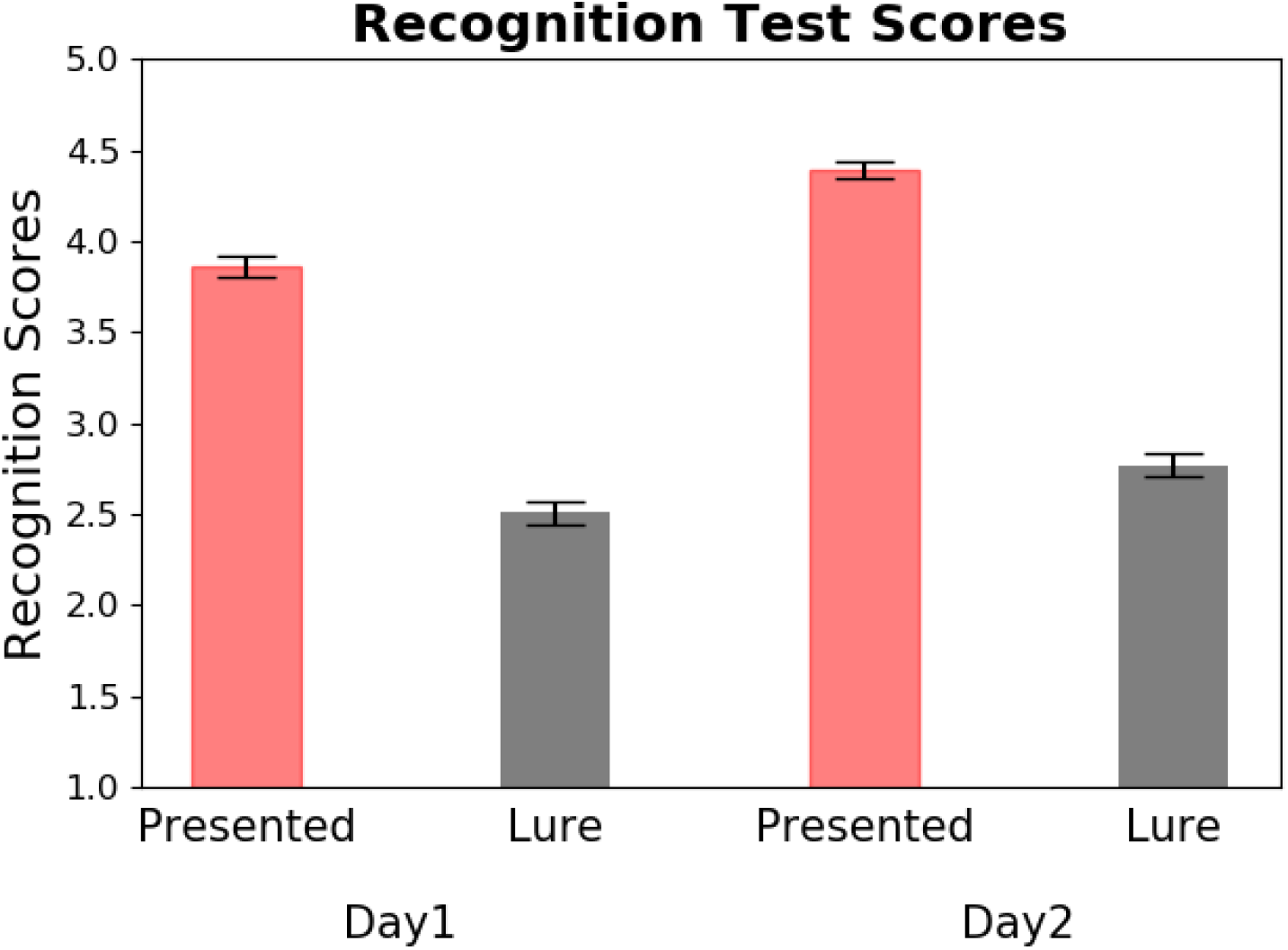
Recognition test scores for both prescan days. Plot shows that presented excerpts were given higher recognition scores than lures. The y-axis represents a 5-point Likert scale where one means not studied and five means studied. Error bars represent standard error of the mean.

On the third day, participants returned for the scanning session in which they listened to the playlist twice (with excerpts played in a different order for the two scanning runs; as noted above, the order of excerpts within a run was the same across participants). During each run, participants were asked to perform a white noise detection task. Specifically, during each excerpt, a brief (1 sec) white noise pulse was played at a randomly chosen time point within the middle 60% of each excerpt. The onset of each noise pulse was also randomized across participants. Participants were told to make a button response to indicate that they heard the noise. This manipulation served to keep participants attentive throughout each excerpt. Following both scanning runs, participants took a final recognition test and then completed a brief demographic survey.

### Event annotations by human observers

In a separate behavioral experiment, we asked seven different raters (only one rater reported having extensive musical training) to listen to our stimuli one at a time with the task of pressing a button when a “meaningful” transition occurred within each piece (similar to the method used by Sridharan et al., 2007). The number of event boundaries identified by the observers varied across excerpts ranging from 3 to 17 boundaries (with a mean of 7.06 and standard deviation of 0.91 across excerpts). It is worth noting that excerpt durations also varied, with a range of 90 sec to 225 sec (durations were either 90, 135, 190, or 225 sec) and an average duration of 157.5 sec and standard deviation of 50.3 sec across excerpts. A timepoint was considered to be an event boundary when at least five annotators marked a boundary within three seconds before or after a given timepoint (method used from Baldassano et al., 2017). The mean number of consensus boundaries across excerpts acquired using this method roughly matched the mean number of boundaries assigned by individual participants across all of the excerpts (with a mean of 7.98 and standard deviation of 2.98 across excerpts).

### Scanning parameters and preprocessing

Imaging data were acquired on a 3T full-body scanner (Siemens Prisma) with a 64-channel head coil. Data were collected using a multi-band accelerated T2-weighted echo-planar imaging (EPI) sequence (release R015) provided by a C2P agreement with University of Minnesota (Auerbach et al., 2013; Cauley et al., 2014; Moeller et al., 2010; Setsompop et al., 2012; Sotiropoulos et al., 2013; Xu et al., 2013): 72 interleaved transverse slices; in-plane resolution = 2.0 mm; slice thickness = 2.0 mm with no inter-slice gap; field of view (FOV) = 208 mm; base resolution = 104; repetition time (TR) = 1000 ms; echo time (TE) = 37 ms; flip angle (FA) = 60 deg; phase-encoding (PE) direction = anterior to posterior; multi-band acceleration factor = 8. Three spin-echo volume pairs were acquired matching the BOLD EPI slice prescription and resolution in opposing PE directions (anterior to posterior and posterior to anterior) for susceptibility distortion correction: TR/TE = 8000/66.60 ms; FA/refocus FA = 90/180 deg; TA = 32s (Andersson et al., 2003).

Additionally, a whole-brain T1-weighted volume was collected: 3D magnetization-prepared rapid gradient echo (MPRAGE) sequence; 176 sagittal slices; 1.0 mm^3^ resolution; FOV = 256 mm; base resolution = 256; TR/TE = 2300/2.88 ms; inversion time (TI) = 900 ms; FA = 9 deg.; PE dir = anterior to posterior; IPAT mode = GRAPPA 2X; TA= 5 min 20 sec.

The EPI volumes were realigned using a 6 parameter rigid-body registration (MCFLIRT; Jenkinson et al., 2002). Given the short effective TR (repetition time) of 1s, slice time correction was not performed. Susceptibility-induced distortions were modeled in the opposing spin-echo volume pairs using the FSL ‘topup’ tool, and the resulting off-resonance field output was provided as input to distortion correct the time series of fMRI data using the FSL ‘applywarp’ tool (Andersson et al., 2003). The susceptibility distortion correction and realignment were applied in a single interpolation step to minimize blurring. Remaining pre-processing and co-registration steps were performed using FEAT (Woolrich et al., 2001, 2004). This included linear detrending, high-pass filtering (330 s cutoff), and spatial normalization to the MNI152 template released with FSL.

### Whole-brain searchlight procedure

We conducted our primary analysis using a whole-brain searchlight approach (Figure 3A). First, all participants’ volumetric data were averaged together and divided into overlapping spherical searchlights, each with a radius of 10 voxels and a stride of 5 voxels (Figure 3B). This resulted in 2,483 searchlights that spanned the whole cortex in MNI space. Only searchlights containing at least 30 voxels were included in the analysis and the mean number of voxels per searchlight was 381.76 voxels with a standard deviation of 168.09 voxels. We assigned the output value for a given searchlight to all voxels within a 5-voxel radius to account for the stride and then averaged the values for voxels where overlap occurred. All analyses below were run separately within each searchlight.

**Figure 3.**
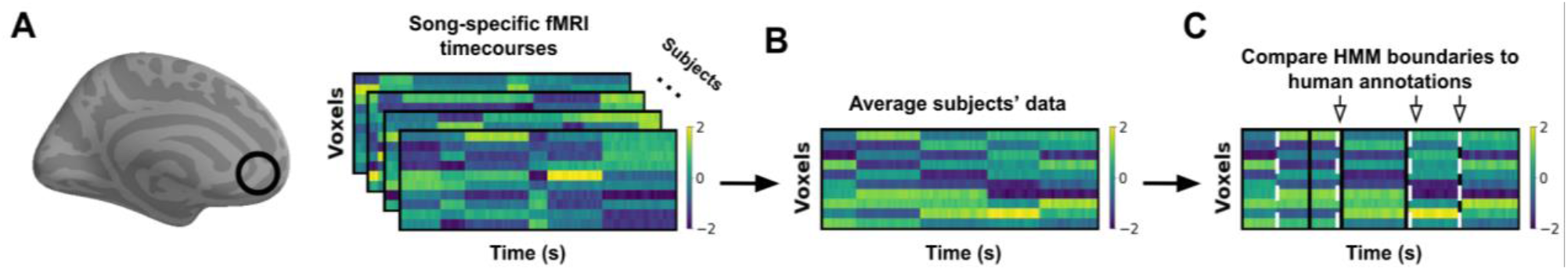
Diagram of analysis pipeline. (From left to right) **A)** For each participant (N=25), voxels from an ROI were selected using a searchlight approach; we then extracted song-specific timecourses (Voxels x TRs (TR=1 second) from the selected voxels (black circle). Inflated brain image was created using PySurfer (https://github.com/nipy/PySurfer/). **B)** Response timecourses were averaged across participants (aligned in volumetric MNI space). **C)** An HMM was used to identify boundary timepoints, when there was a change in the spatial pattern of activity across voxels. HMM boundaries (white dashed lines) and human annotations (black lines) were considered to match (downward arrows) when HMM boundaries fell within 3 TRs (3 seconds) of a human annotation. Then, true match scores were compared to a null distribution constructed by comparing shuffled HMM boundaries to human annotations, resulting in a z-score for each ROI.

### Event segmentation analysis

For each searchlight, we fit a hidden Markov model (Baldassano et al., 2017) to the response time series for each excerpt, setting the number of states in the HMM equal to the number of segments specified by our human annotators for each excerpt. Furthermore, although we provide the HMM with a specific number of events, we do not give it any information about where these events are in the data. Therefore, the model is unsupervised in terms of locating the boundaries between events. We used a specialized HMM variant developed by Baldassano et al. (2017) that is optimized for event segmentation (i.e., identifying jumps in neural patterns). This HMM variant seeks to model the fMRI time series as a set of successive transitions between stable states, where – in our variant of the HMM – the model is not permitted to return to a state once it leaves that state. Fitting the model to the data involves estimating the voxel pattern for each stable event state as well as the timing of transitions between these patterns; this HMM variant was implemented using the EventSegment function in BrainIAK (Kumar et al., 2020).

For our primary analysis, we were interested in finding brain regions whose transition structure most closely resembled the event boundary structure given by our annotators (Figure 3C). After acquiring boundary estimates from the HMM, we evaluated how closely in time the boundaries found by the model matched the boundaries supplied by our annotators. To quantify the degree of match, we counted the number of human-annotated boundaries for which there was an HMM boundary within three TRs (3 seconds) of that human-annotated boundary. Note that all human boundaries were shifted forward 5 TRs (5 seconds) to account for the hemodynamic lag. We created a null model by randomly selecting timepoints as boundaries (keeping the number of events the same, as in Baldassano et al. (2017) and computed the number of matches for these null boundaries, repeating this process 1000 times to produce a null distribution. We computed a z-value of the real result versus the null distribution by subtracting the average of the permuted match scores from the true match score and dividing this difference by the standard deviation of the permuted scores. This procedure was repeated at every searchlight. By acquiring z-scores at each searchlight for all 32 excerpts (16 distinct excerpts x 2 runs), we obtained 32 separate spatial maps of z-scores. Next, we averaged the two z-maps corresponding to each distinct excerpt (one from each run), resulting in 16 total z-maps. To summarize across the z-scores for the 16 distinct excerpts, we ran a one sample t-test against zero to see which voxels had the most reliable matches across all excerpts. The resulting t-values were converted to p-values and then adjusted for multiple tests to control for the false discovery rate (FDR) at a value q (Benjamini & Hochberg, 1995).

To visualize the results, each spatial map of t-values was displayed on the cortical surface (masked to include only vertices that exhibited a significant effect). Since each analysis was performed in volumetric space, volume data were projected to the cortical surface using the automatic volume to surface rendering algorithm within PySurfer (https://github.com/nipy/PySurfer/).

### Controlling for acoustic features

To further determine whether regions of the DMN represent high-level musical event structure, as opposed to surface-level acoustic information, we repeated the searchlight analysis, this time regressing out musical features extracted from each auditory stimulus prior to fitting the HMM. All feature extraction was performed using Librosa (McFee et al., 2015), a Python package developed for audio and music analysis. These features consisted of mel-frequency cepstral components (MFCCs; i.e. timbre information), chromagrams (tonal information), tempograms (rhythmic information), and spectrograms. For MFCCs, the top 12 channels were extracted since these lower-order coefficients contain most of the information about the overall spectral shape of the source-filter transfer function (Poorjam, A.H., 2018). Chromagrams consisted of 12 features, each corresponding to a distinct key in the chromatic scale. Tempograms initially consisted of 383 features, each representing the prevalence of certain tempi (in beats per minute; BPM) at each moment in time. Since most of the tempo-related variance was explained by a much smaller set of features, we reduced the 383 features to 12 features using PCA (variance explained was 99%) in order to match the number of features used for MFCCs and chromagrams. Spectrograms were extracted using the short-time Fourier transform (STFT) and then converted to a dB-scaled spectrogram. Then, we also used PCA to reduce the dimensionality of the spectrograms to 12 components, which explained 98% of the frequency-related variance. For the final step of this analysis, we applied the HMM to the residuals after the musical features were regressed out of the neural data.

### Identifying preferred event timescales

After identifying brain regions with neural event boundaries that matched the human annotations (using the procedures described in *Event segmentation analysis*, above), we ran a follow-up analysis to further probe the properties of four such regions (bilateral auditory cortex, bilateral angular gyrus, bilateral mPFC, and bilateral precuneus). Specifically, the goal of this follow-up analysis was to assess the preferred timescales of these regions. Angular gyrus, mPFC, and precuneus were selected (in addition to auditory cortex) because activity patterns in these regions have been found to exhibit high-level event structure in recent studies using naturalistic stimuli such as movies (Baldassano et al., 2017; Ben-Yakov & Henson, 2018; Geerligs et al., 2021; Honey et al., 2012) and spoken narratives (Lerner et al., 2011). In contrast to our primary event segmentation analysis (which used a fixed number of events for each excerpt, matching the number of human-annotated events for that excerpt), here we tried models with different numbers of events and assessed how well the model fit varied as a function of the number of events. The measure of model fit we used was the average pattern similarity between pairs of timepoint-specific multivoxel patterns falling *within* the same event, minus the average pattern similarity between patterns falling *across* events (Baldassano et al., 2017). We call this measure the *WvA score* (short for “Within versus Across”); higher WvA scores indicate a better fit of the event boundaries to the data. The ROIs for this analysis were defined by selecting voxels within functionally-defined parcellations (Schaefer et al., 2018) corresponding to bilateral auditory cortex, bilateral angular gyrus, bilateral mPFC, and bilateral precuneus, and then (for extra precision) intersecting these parcels with voxels that were also significant in our primary searchlight analysis looking for neural boundaries that matched human-annotated boundaries (q < 0.01). For each ROI, we fit HMMs to each song with differing numbers of events ranging from 3 to 45. For each HMM fit, we measured the maximum event duration, and then identified all pairs of timepoints whose temporal distance was less than this duration. The constraint of using timepoints whose distance was less than the maximum event duration was used so that the number of within-and across-event pairs would be roughly equal (regardless of the number of events). The WvA score was computed as the average spatial pattern correlation for pairs of timepoints falling in the same (HMM-derived) event minus the average correlation for pairs of timepoints falling in different events. We then averaged the results across excerpts. Note that, since the excerpts are different lengths, a given number of events might correspond to different average event lengths for different excerpts (e.g., a 3-event model applied to a 180 second excerpt has an average event length of 60 seconds, but a 3-event model applied to a 90 second excerpt would have an average event length of 30 seconds). Because our goal was to find each area’s preferred event length, we converted our WvA results for each excerpt to be a function of the average event length (in seconds) rather than the number of events, and averaged these results across excerpts. Finally, to compute the preferred event length for each ROI, we identified the range of event lengths that were within 5% of the maximum WvA score for that ROI; we report the midpoint of this range as the preferred event length.

To test whether the preferred event length in auditory cortex was shorter than that of angular gyrus, precuneus, and mPFC, we performed a bootstrap analysis, repeating the above analysis (including re-running SRM) 1000 times for different bootstrap resamples of the original dataset. At each iteration of the bootstrap, we applied the analysis to a sample of participants drawn randomly with replacement from the original data. We computed p-values by finding the proportion of bootstraps where the preferred length for auditory cortex was greater than the preferred length for angular gyrus, precuneus, and mPFC.

## Results

### Neural boundary match to behavioral annotations

We wanted to test the hypothesis that behaviorally-defined event boundaries could be identified in higher-order cortical regions, especially those overlapping with the DMN. For this analysis, we fit an HMM to BOLD data averaged across both runs and then compared the HMM boundaries to the specific timepoints labeled as boundaries by the annotators. We found significant matches between model boundaries and human annotations in auditory cortex, angular gyrus, precuneus, and medial prefrontal cortex. (Figure 4). Results for this analysis are split by run 1 and run 2 in Appendix B.

**Figure 4.**
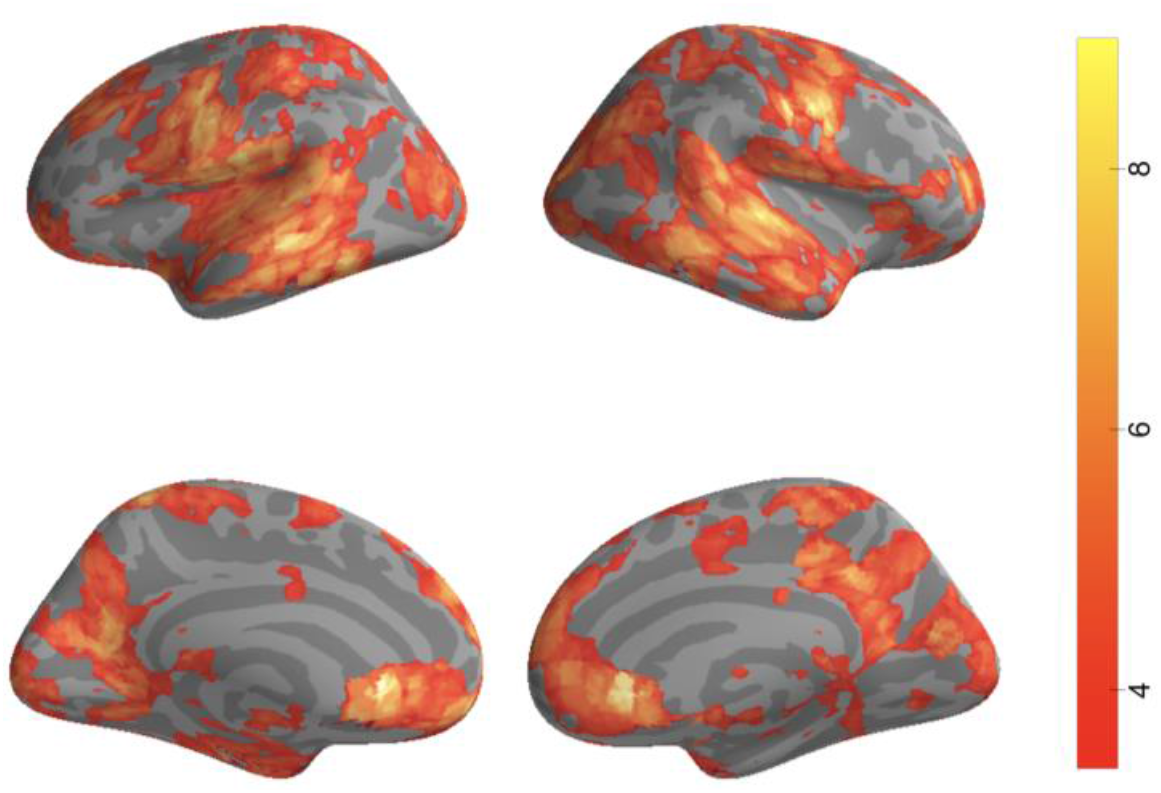
Distance to Boundary Searchlight Results. For 2,483 searchlights spanning the entire cortex, we tested whether the average match between neural and annotated boundaries across all songs was significantly greater than zero. Significant voxels overlapped with auditory cortex as well as areas of the default mode network such as precuneus, medial prefrontal cortex, and angular gyrus. Results are thresholded via FDR, q < 0.01.

### Influence of acoustic features

To determine the extent to which the neural event boundaries were driven by acoustic features, we also performed a version of the searchlight analysis in which we controlled for spectral, timbral, harmonic, and rhythmic information. Overall, this reduced the number of searchlights passing the q < 0.01 false discovery rate threshold (Figure 5) compared to the original searchlight analysis. However, searchlights in DMN regions (precuneus, angular gyrus, and mPFC) did pass the q < 0.01 threshold, with voxels in mPFC being (numerically) least affected by the feature removal. When we set a more liberal false discovery rate threshold (q<0.05; results shown in Appendix A), the relationship between neural event boundaries and human annotations was still largely conserved in precuneus, angular gyrus, and auditory cortex. This suggests that, although voxels in precuneus and angular gyrus are more sensitive to acoustic features than mPFC, event boundaries found in these regions do not directly correspond to simple changes in the acoustic features and may instead be related to more complex representations of the event structure (e.g. non-linear combination of acoustic features). Notably, significant searchlights in auditory cortex were also observed (particular in right auditory cortex), indicating that – even in sensory areas – the event boundaries were being driven (at least in part) by more high-level aspects of the music.

**Figure 5.**
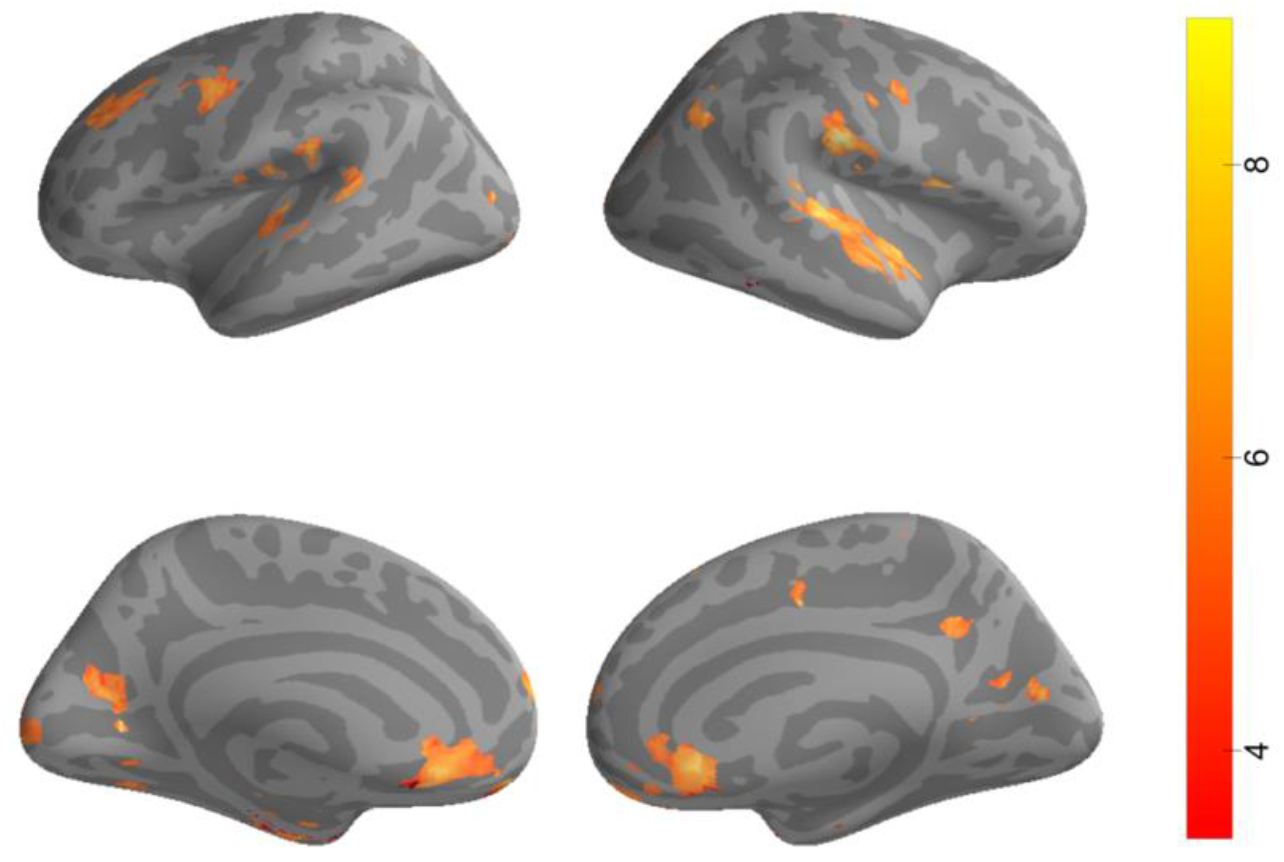
Searchlight results accounting for acoustic features. We recomputed the match between HMM-derived neural boundaries and human annotations after regressing out acoustic features from each participant’s BOLD data prior to fitting the HMM. Significant effects were still observed in parts of the DMN as well as auditory cortex, suggesting that boundaries detected in these areas do not necessarily depend on acoustic features. Results are thresholded via FDR (q < 0.01).

### Comparing annotated event boundaries to changes in acoustic features

In a follow-up analysis, we sought to further investigate the relationship between the event boundaries and changes in the acoustic features by assessing how often the behaviorally-defined event boundaries occurred at the same time as changes in each of the acoustic features. In other words, how often does a change in an acoustic feature generate a human boundary? To estimate the number and locations of state changes within each of the excerpts, we applied the Greedy State Boundary Search event segmentation model (GSBS; Geerligs, van Gerven, & Güçlü, 2021) to each of the acoustic features (i.e. MFCC, chromagram, tempogram, spectrogram) extracted from each of the excerpt audio files. One advantage of using the GSBS algorithm for this analysis is that GSBS can automatically identify the optimal number of states that maximizes the difference between within-vs. across-event similarity. After acquiring the optimal set of GSBS event boundaries for each excerpt, we compared them to the human annotations by computing the probability that a shift in an acoustic feature generated a matching human annotation (within 3 seconds). Additionally, we assessed whether this probability was greater than what would be predicted by chance by establishing a null distribution whereby we shuffled the feature boundaries for each excerpt while preserving the distances between boundaries. We found that feature boundaries did align with human-annotated boundaries more often than in the null model, but that most feature changes did not result in a human-annotated boundary (p(annotation | chroma boundary) = 0.143, vs null value of 0.115 [p<0.001]; p(annotation | MFCC boundary) = 0.493, vs null value of 0.299 [p<0.001]; p(annotation | tempo boundary) = 0.198, vs null value of 0.179 [p<0.05]; p(annotation | spectrogram boundary) = 0.160, vs null value of 0.128 [p<0.001]) (illustrated using an example excerpt in Figure 6A).

**Figure 6.**
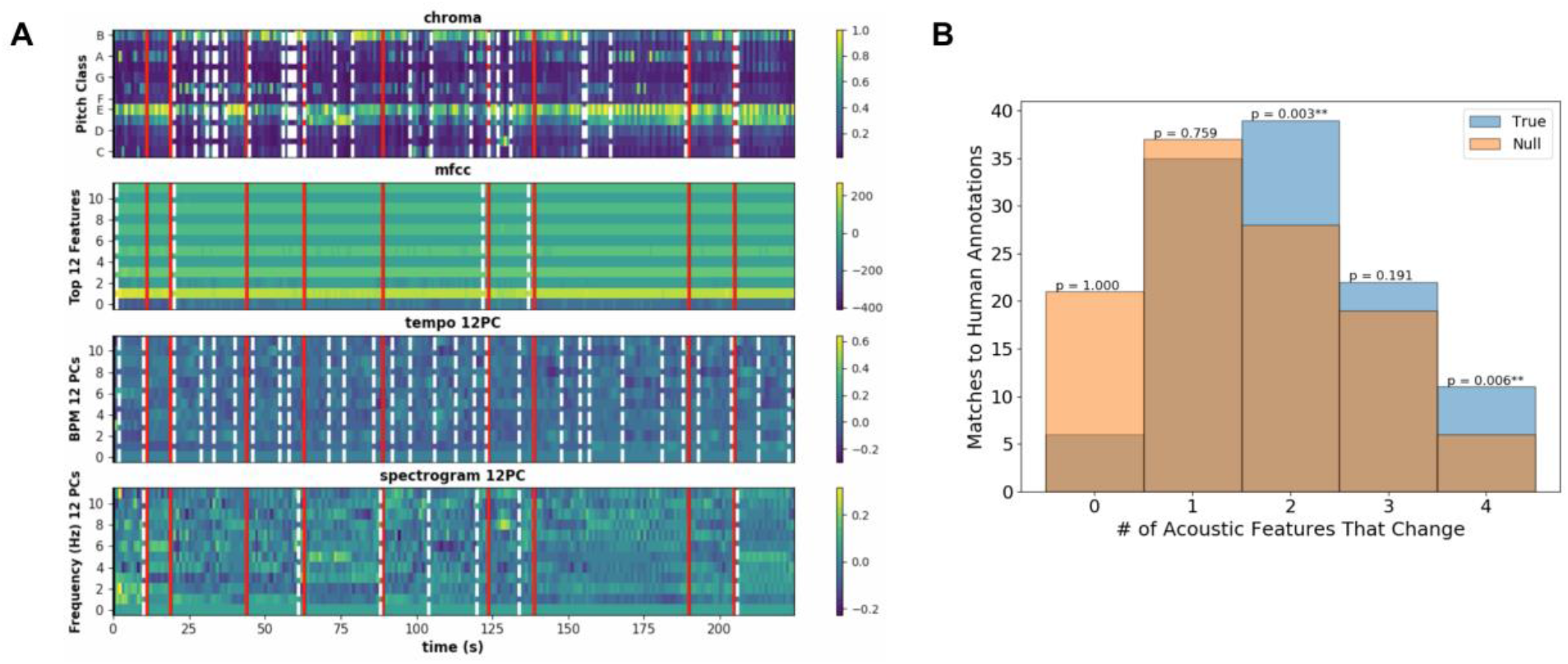
Panel A. Example acoustic features (from *My Favorite Things* by John Coltrane) showing overlap between human annotations (red) and feature boundaries (white dashed lines). For each acoustic feature, we identified timepoints at which changes occurred using the GSBS event segmentation model (white dashed lines). We then compared the locations of these feature boundaries to the locations of the human annotations (red lines); see text for results. **Panel B. Number of acoustic features that change at human-annotated event boundaries**. Counting how many acoustic features exhibit a boundary at the same time as a human-annotated boundary (blue) versus a null distribution (orange), we find that the observed distribution is shifted upward relative to the null distribution, such that human-annotated boundaries are more likely to occur in response to two or more feature changes. Furthermore, some human annotations occur in the absence of any feature change.

We also computed the distribution (across human-annotated boundaries) of the number of acoustic feature types that changed within 3 seconds of each annotated boundary (e.g., if chroma and tempo both changed, that would be two feature types). We compared this distribution to a null model that we obtained by shuffling the human-annotated boundaries for each excerpt while preserving the distances between boundaries. The results of this analysis are shown in Figure 6B. The fact that the observed distribution is shifted upward relative to the null tells us that probability of human boundaries coinciding with auditory feature changes is higher than would be expected due to chance (*X*^2^ = 19.54, p<0.001 by permutation test). The figure also shows that – while the majority of human boundaries occurred at points where 2 or more acoustic feature changes were present – some human boundaries occurred at time points where no acoustic feature changes were present.

### Preferred event lengths across ROIs

How do we reconcile the role of auditory cortex in high-level event representation (as shown in the above analyses) with its well-known role in representing low-level auditory features? Importantly, these claims are not mutually exclusive. Our analyses, which set the number of event states in the model to equal the number of human-annotated boundaries, show that auditory cortex has some (statistically-reliable) sensitivity to high-level events, but this does not mean that this is the *only* event information coded in auditory cortex or that it is the *preferred* level of event representation.

We defined the preferred timescale of each region (ROI selection is discussed in the Experimental design and statistical analysis section) by running HMMs with different numbers of event states, and finding the average event length (in seconds) that produced the best model fits across songs (Figure 7A). Using a bootstrap analysis, we found that auditory cortex’s preferred event length (13.81 s) was significantly shorter than the preferred event length of mPFC (25.59 s; p=0.009) but was not significantly shorter than the preferred length of angular gyrus (13.36 s; p=0.664) or precuneus (14.61s; p=0.338). The preferred event length in mPFC was also significantly longer than the preferred event length for precuneus (p=0.017) and angular gyrus (p=0.004).

**Figure 7.**
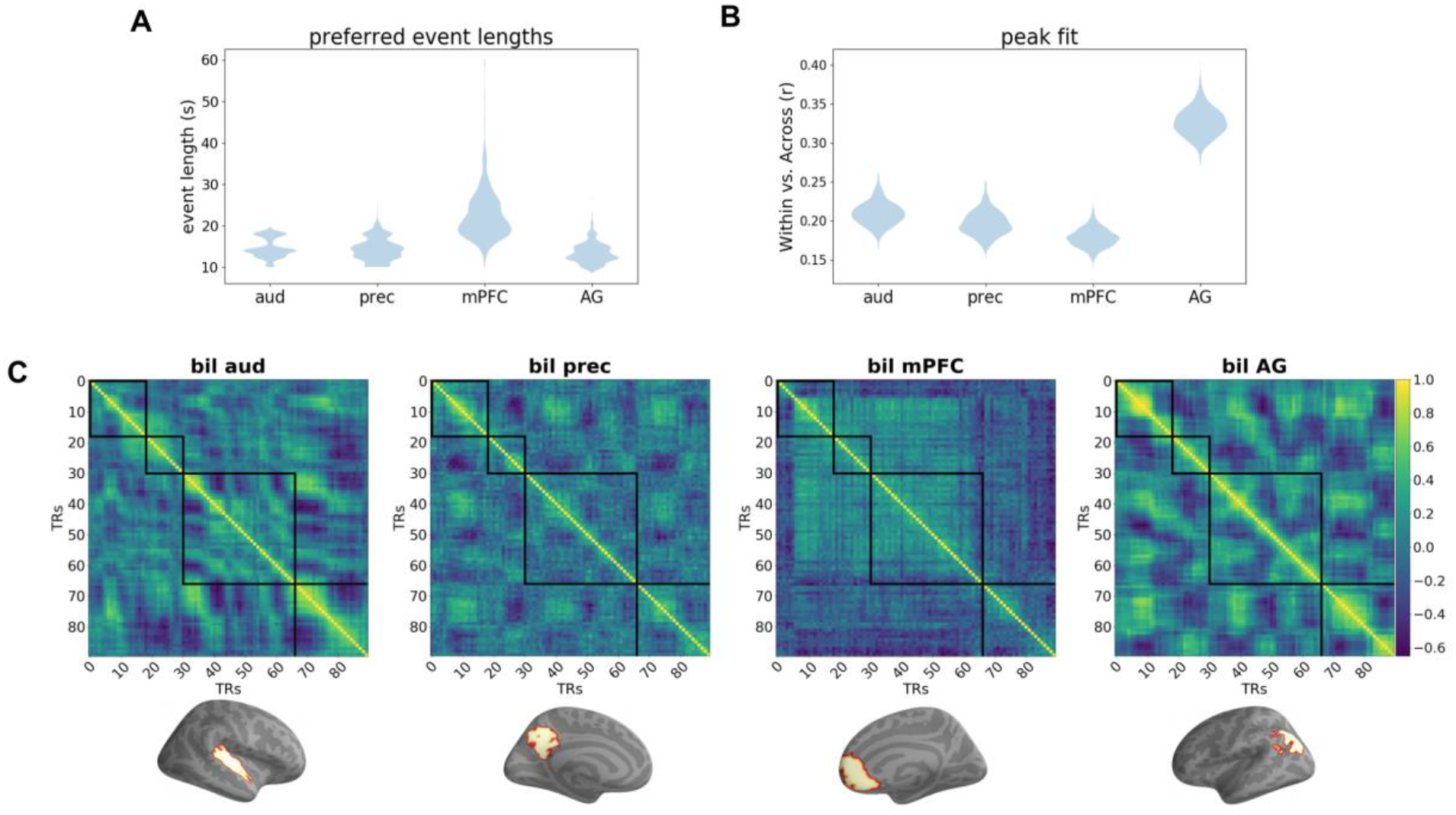
**Panel A**. Longer states were preferred in mFPC (average event length 25.59 s) than in auditory cortex (13.81 s), precuneus (14.61 s), and angular gyrus (13.36 s). The preferred event length did not significantly differ between auditory cortex, precuneus, and angular gyrus. **Panel B**. The overall within-event pattern similarity was highest in angular gyrus, suggesting that the stability of musical event representations was higher than in other ROIs. There was no difference in within-event pattern similarity between precuneus and auditory cortex, however, pattern similarity was significantly less in mPFC than in auditory cortex (p < 0.05). **Panel C**. Similarity matrices (for the first 90 seconds of the excerpt *Capriccio Espagnole* by Nikolai Rimsky-Korsakov) showing pattern similarity over time for each region of interest. mPFC exhibits the largest event structure relative to auditory cortex, precuneus, and angular gyrus.

In addition to varying in timescale (i.e., in the best-fitting number of events), regions could differ in the quality of this fit; some regions may exhibit sharper event transitions, with large pattern changes across HMM event boundaries and highly stable patterns within events. We therefore tested whether the model fit (maximum WvA score) was different between the four ROIs (Figure 7B). We found that the model fit for angular gyrus was significantly greater than auditory cortex (p<0.001), precuneus (p<0.001), and mPFC (p<0.001), indicating that the temporal event structure was strongest in angular gyrus. For analyses of preferred event length and model fit in a more complete set of DMN ROIs and in hippocampus, see Appendices C and D, respectively.

## Discussion

In this study, we sought to determine whether brain areas that have been implicated in representing high-level event structure for narrative-based stimuli, such as movies and spoken narratives, are also involved in representing the high-level event structure of music in a group of passive listeners. We provide evidence that regions of the default mode network (DMN) are involved in representing the event structure of music as characterized by human annotators. The durations of these human-annotated events lasted on the order of a few seconds up to over a minute.

Our results indicate that high-level structure is represented in both high-level DMN regions but also in auditory cortex. Auditory cortex, however, may not explicitly represent high-level events at the level of human annotators; that is, the behaviorally-identified event boundaries are likely a subset of the finer-grained event boundaries encoded in auditory cortex. When we force the HMM to match the number of human-annotated boundaries, the HMM finds them, demonstrating that coding in auditory cortex is modulated by high-level event structure. However, when we remove this constraint and allow the number of events to vary, auditory cortex prefers shorter events on average relative to mPFC, but not precuneus and angular gyrus (Figure 7 Panel A), whereas mPFC preferred the longest events compared to the other three ROIs. The finding that the preferred event length of auditory cortex was not significantly different from that of precuneus and angular gyrus was surprising given the prediction that auditory cortex, which is generally thought to respond to fast-changing aspects of a stimulus, would represent shorter events than higher-order brain areas (Baldassano et al., 2017; Farbood et al., 2015; U. Hasson et al., 2008; Lerner et al., 2011); we discuss this point further in the limitations section below. In addition to measuring each area’s preferred timescale, we also measured within-event stability across the four ROIs; here, we found that angular gyrus exhibits the strongest within-event activity relative to precuneus, mPFC, and auditory cortex.

Next, we showed that – when we regress out acoustic features corresponding to timbre, harmony, rhythm, and frequency amplitude and re-run the analysis – voxels in higher-order areas (mPFC, angular gyrus, and precuneus), as well as auditory cortex, still significantly match with the annotations. These results suggest that event boundaries in these regions are not purely driven by acoustic changes in the music, but are also tracking more complex event structure in musical pieces. These findings are consistent with findings from (Abrams et al., 2013), who found that naturalistic music elicited reliable synchronization in auditory cortex as well as higher-order cortical areas after controlling for acoustic features; they concluded that this synchronization was not purely driven by low-level acoustical cues, and that it was likely driven by structural elements of the music that occurred over long timescales.

To further determine how much event boundaries were driven by changes in acoustic features, we ran a follow-up analysis where we first identified event transitions in each of the acoustic features corresponding to timbre, tonality, rhythm, and frequency amplitudes for each excerpt using an unsupervised algorithm (GSBS); then, we computed the probability that a human annotation was generated by changes in each of the different types of acoustic features. We found that the probability of human-annotated boundaries coinciding with acoustic feature changes was higher than the rate expected due to chance, but the relationship was complex: While changes in each of the individual acoustic feature types were significantly related to the occurrence of annotated boundaries, none of these features came close to fully predicting the annotated boundaries; and while the majority of annotated boundaries occurred at time points where two or more acoustic features changed, some annotated boundaries did not correspond to changes in any of the acoustic features that we tracked. This adds further support to the possibility that boundaries marking the shift between large-scale segments within the DMN and auditory areas could be driven by a complex shift in a combination of the acoustic properties and/or possibly emotional (Daly et al., 2015) or narrative (McAuley et al., 2021; Margulis et al., 2019; Margulis et al., 2021) changes within the excerpts, rather than a change in a single feature.

Importantly, our findings of high-level coding in auditory cortex converge with other recent work demonstrating that hierarchical neural representations of music are distributed across primary and non-primary auditory cortex (Landemard et al., 2020) and that higher-order representations of music in these areas may even support complex behaviors such as genre recognition in humans (Kell et al., 2018). Our study contributes to this growing literature by showing that auditory cortex codes for musical event representations at intermediate timescales (∼14s). Note also that auditory cortex coding for these intermediate-scale events is not mutually exclusive with it *also* coding for shorter-timescale events. When discussing limitations of our study below (limitation point number 4), we provide some reasons why our design was not ideal for detecting neural coding of short-timescale events.

In our study, we provide strong evidence for the involvement of medial prefrontal cortex in representing high-level musical event structure. Recent fMRI studies of naturalistic stimulus processing (i.e., audiovisual movies) have shown that mPFC may perform event segmentation and integration during continuous memory formation (Antony et al., 2021; Liu et al., 202) and that events in this region can last up to hundreds of seconds (Hasson et al. 2015; Chen et al. 2017; Geerligs et al. 2021). We also show that the preferred event length in mPFC was ∼25 seconds (which was roughly equal to the preferred timescale found for mPFC in the study by Geerligs et al. (2021) in which a movie was used rather than music), adding further support to the hypothesis that mPFC plays an important role in representing long-timescale information in naturalistic stimuli. Furthermore, our findings go beyond the assumption that areas of the DMN only represent long-timescale information for narrative-based stimuli and instead suggest that areas of the DMN represent long-timescale information across a range of naturalistic stimuli, including music. The recruitment of mPFC during music processing has also been found in a previous study (Blood and Zatorre, 2001). Specifically, Blood and Zatorre showed that activity in vmPFC was correlated with pleasure response ratings to music, suggesting that frontal areas, which represent long-timescale event structure for music, may also play a role in processing reward and affect in response to music.

Our findings that precuneus, mPFC, and angular gyrus were involved in representing high-level musical event structure contrast with those in Farbood et al. (2015), who found that regions that responded reliably to stories did not respond reliably to music. Furthermore, in their study, there was minimal overlap between voxels in angular gyrus and mPFC that responded to stories and voxels that responded to music. In our study, we show that, at a regional level, these areas are indeed involved in representing the high-level event structure in music. One major way in which our studies differed was our use of an HMM to detect evidence of musical event structure in higher-order areas. The HMM is optimized to detect periods of relative stability punctuated by shifts in response patterns, which one would expect for an area encoding high-level event structure (i.e., there should be stability within events and changes across events). Temporal ISC (inter-subject correlation analysis; the analysis method used in the study by Farbood) is designed to pick up on *any* kind of reliable temporal structure and is not specifically designed to detect the “stability punctuated by shifts” structure that we associate with event cognition, making it less sensitive to this kind of structure when it is present. This highlights one of the advantages of using HMMs for detecting meaningful brain activity related to the temporal dynamics of naturalistic stimuli, such as music.

Our study had several limitations. First, in our feature regression analysis, the acoustic features we selected may not represent the full range of acoustic dynamics occurring throughout each excerpt. Previous studies using encoding models to examine brain activity evoked by music employed a range of acoustic features, such as the modulation transfer function (Norman-Haignere et al., 2015; Patil et al., 2012) as well as music-related models representing mode, roughness, root mean square energy (RMS), and pulse clarity (Alluri et al., 2012; Nakai et al., 2021; Toiviainen et al., 2014). However, the types of information captured by these features are also roughly captured by the features used in this study. For example, features representing roughness and RMS capture timbral information while pulse clarity captures rhythmic information. On the other hand, although these features capture some information related to the ones used in this study they may nonetheless still be useful for capturing additional information not fully captured by our features. Future work is needed to determine how higher-order areas are affected by a larger set of acoustic features.

(2) Another caveat is that we only scanned participants listening to pre-familiarized musical stimuli – as such, it is unclear whether the observed pattern of DMN results (showing engagement of these regions in long-timescale segmentation) would extend to unfamiliar musical stimuli. Consistent with this view, work by Castro et al. (2020) showed that familiar music engaged DMN more strongly than unfamiliar music. However, a study by Taruffi et al. (2017) showed that DMN was engaged for unfamiliar music, particularly for sad music compared to happy music. Future work investigating high-level musical event structure representation can address this by scanning participants while they listen to both unfamiliar and familiar stimuli.

(3) The white-noise-detection task that participants performed may have influenced DMN responding. The DMN has been shown to activate during mind-wandering or stimulus independent thought (SIT; Mason et al., 2007). Since the white noise was spectrally distinct from the music, participants could conceivably perform the white-noise-detection task without attending to the music, leaving room for them to mind-wander in between white noise bursts; consequently, some of the DMN responding could (in principle) have been driven by mind-wandering instead of music-listening. However, stimulus-independent mind-wandering cannot explain our key finding that neural event boundaries in DMN regions align with the annotated event boundaries – this result clearly demonstrates that these DMN areas are tracking structural aspects of the music.

(4) It is possible that our estimates of preferred event length for different ROIs were biased by the range of event lengths present in our stimulus set. In particular, a lack of short (vs. long) events may have resulted in an upward bias in our estimates of preferred event length. This bias, however, cannot explain the relative differences that we observed between ROIs’ preferred timescales, such as mPFC preferring longer events than auditory cortex, precuneus, and angular gyrus. However, the relative scarcity of short events may have impaired our ability to resolve timescale differences between regions at the short end of the timescale continuum; in particular, this might help to explain why we did not observe significant differences in preferred timescales between primary auditory cortex (which, based on prior work, we expected to have a short timescale preference) and DMN regions. Future work can shed light on this by using stimuli with a broader range of event lengths. However, even if we include stimuli with shorter events, our ability to detect these more rapid event transitions may be inherently limited by the slow speed of the fMRI BOLD response.

### Conclusion

In this study, we sought to determine whether certain regions in the default mode network, which have been shown to be involved in representing the high-level event structure in narratives, were also involved in representing the high-level event structure in real-world music. Recent fMRI work, not using music, has shown that hidden Markov models (HMMs) can help us understand how the brain represents large-scale event structure. By using HMMs to segment fMRI response patterns over time according to the event structure provided by a separate group of human annotators, we found that areas of the DMN were indeed involved in representing the high-level event structure (e.g., phrases, sections) in music in a group of passive listeners. Of particular importance are the findings that mPFC has a chunking response that is close to that of human observers and survives the boundary alignment searchlight analysis even after controlling for acoustic features. This suggests that mPFC plays an important role in high-level event representation not only for movies and stories (Lerner et al., 2011; Hasson et al. 2015; Chen et al. 2017; Baldassano et al., 2017; Geerligs et al. 2021) but also for instrumental music.

## Appendix A

**Appendix A.**
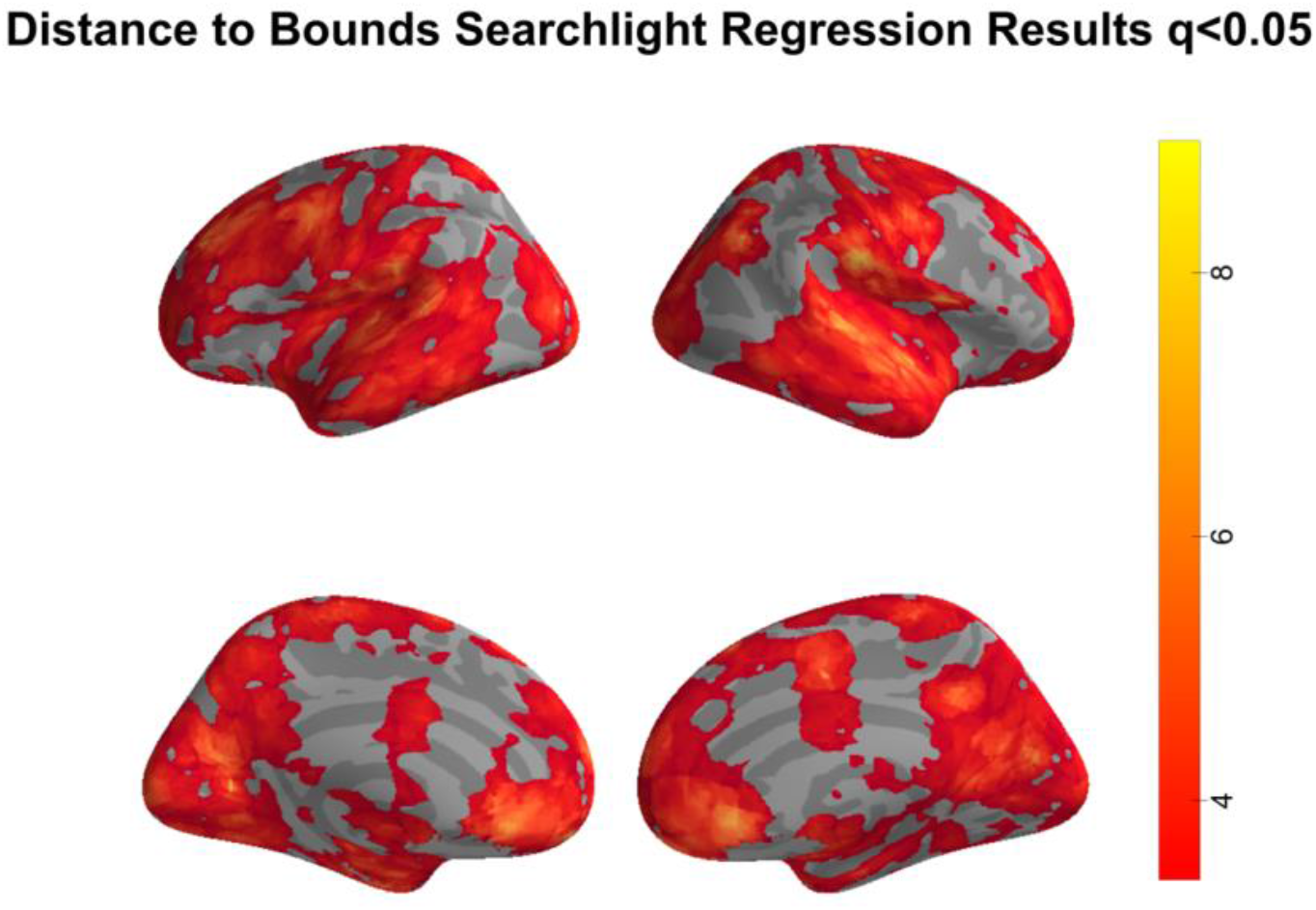
Distance to Boundary Regression Results at q<0.05. Plots show distance to boundary regression results in which we regress out MFCCs, chromagrams, tempograms, and spectrograms. Results are FDR-corrected at q<0.05. These results show that, although many voxels in the DMN are not significant at the q<0.01 threshold (Figure 5), many DMN voxels do survive when we threshold the regression results at q<0.05. This suggests that, while many voxels in the DMN are somewhat sensitive to acoustic features (since many of these voxels do not survive at q<0.01 in the non-regression distance to boundary results), activity in these areas is not solely driven by low-level acoustic features.

## Appendix B

**Appendix B.**
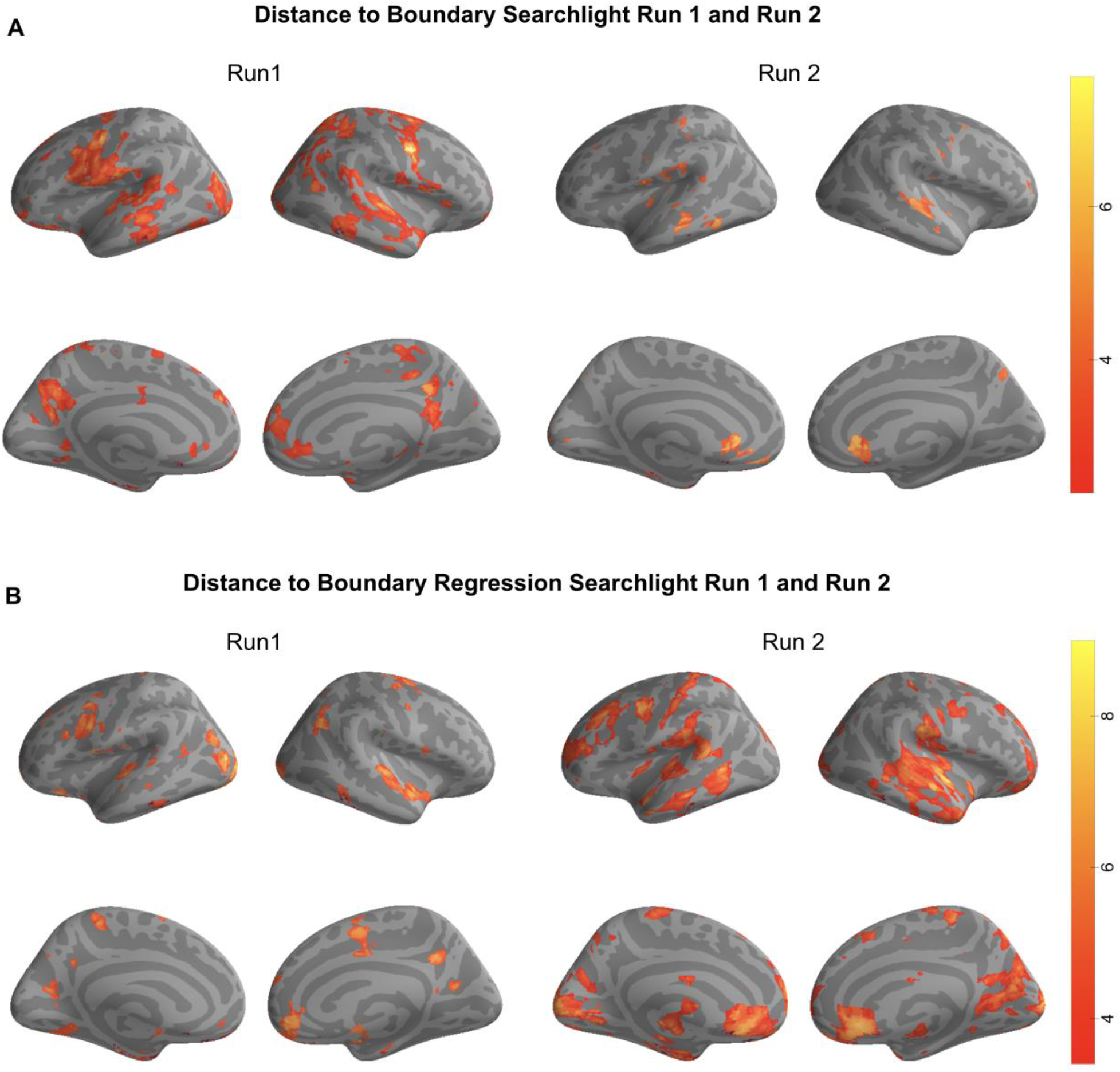
Panel A. Distance to Boundary Searchlight Run 1 and Run 2. Searchlight maps for each run separately showing regions where significant matches between human annotations and HMM boundaries were observed (FDR corrected q < 0.01). **Panel B. Distance to Boundary Regression Searchlight Run 1 and Run 2**. Searchlight maps for each run separately showing regions where significant matches between human annotations and HMM boundaries were observed after regressing out acoustic features (FDR corrected q < 0.01).

## Appendix C

**Appendix C.**
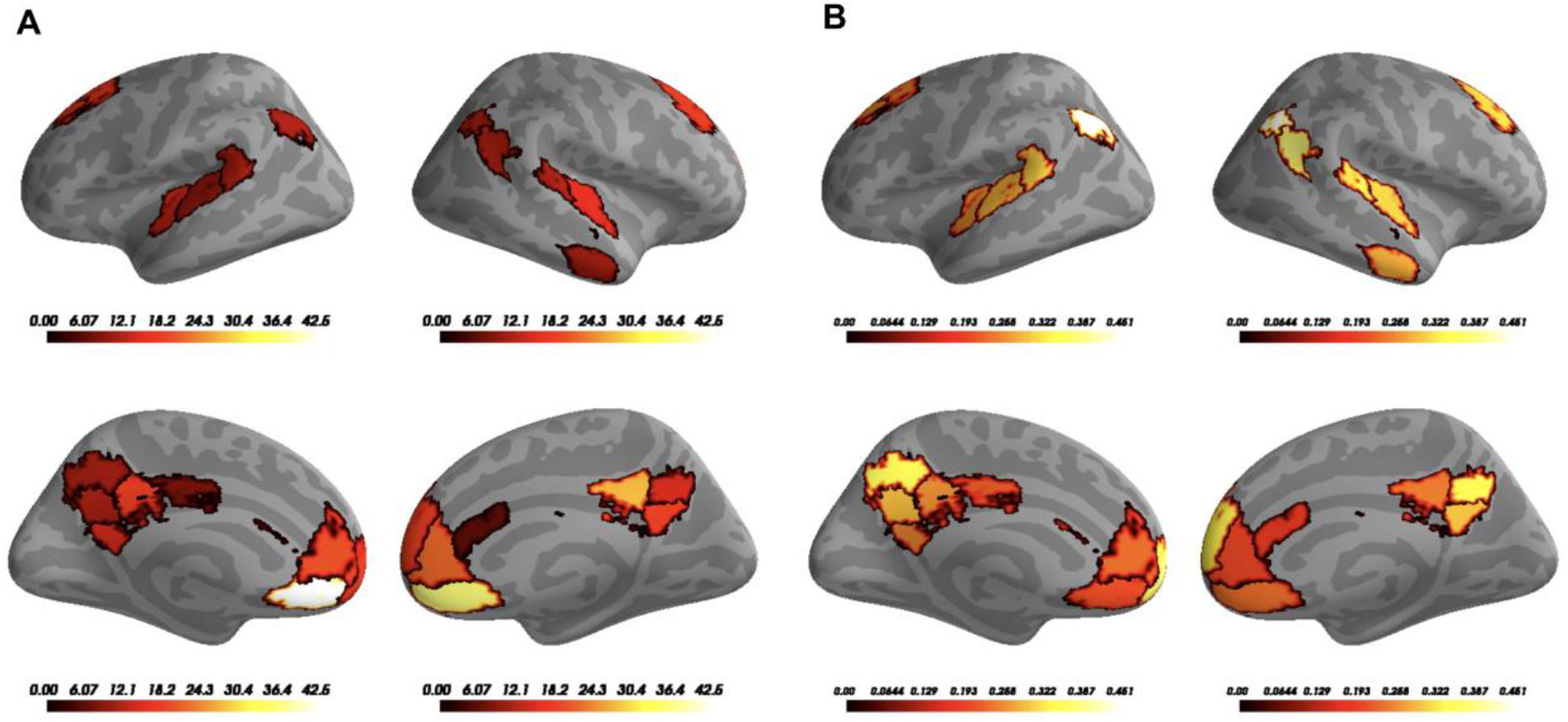
**Panel A**. Preferred event lengths across finer set of DMN and auditory ROIs. We sought to further determine the set of event lengths preferred within each ROI using a finer set of parcellations (Schaefer 300 as opposed to Schaefer 100). We attempted to threshold this image by only including ROIs with significant model fits (determined via bootstrapping). Nothing survived our threshold criteria, therefore we are reporting un-thresholded results. Sub-regions of DMN preferred a variety of event lengths, which was not obvious when using a coarser set of parcellations. For example, although mPFC obtained from the Schaefer 100 parcellation set shows a preference for the longest event lengths (∼25s), when evaluating this for a finer set of mPFC ROIs (Schaefer 300), we can see that mPFC sub-regions prefer a variety of event lengths ranging from 6s-40s. **Panel B**. Model fits also vary greatly for the same set of DMN and auditory ROIs.

## Appendix D

**Appendix D.**
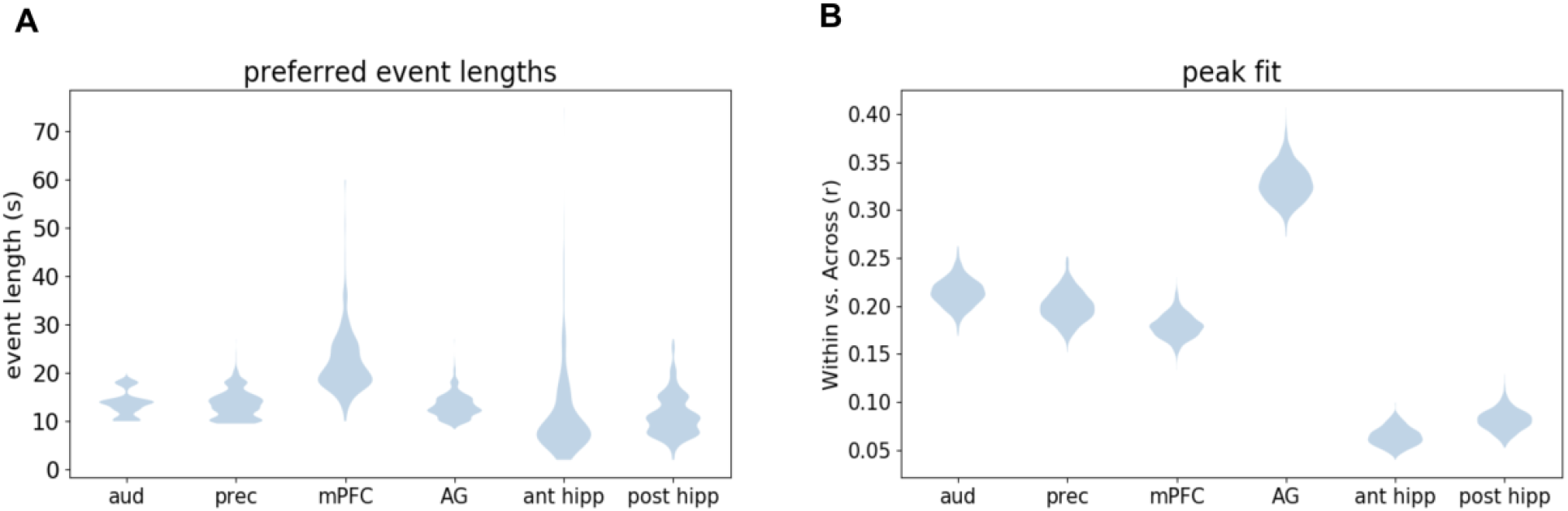
**Panel A**. Anterior hippocampus preferred event length did not significantly differ from auditory cortex, precuneus, mPFC, angular gyrus, or posterior hippocampus. Posterior hippocampus preferred event length did not significantly differ from auditory cortex, precuneus, angular gyrus, or posterior hippocampus, but was significantly less than mPFC (p<0.05). **Panel B**. Our measure of model fit (i.e., the difference between within-event and across-event pattern similarity) was significantly lower in hippocampal ROIs than in other DMN ROIs [auditory cortex (p<0.001), precuneus (p<0.001), mPFC (p<0.001), angular gyrus (p<0.001)] while model fit in posterior hippocampus was greater than in anterior hippocampus (p<0.05).

## Acknowledgments

We thank Mark A. Pinsk for contributing to the *Scanning parameters and preprocessing* section of the manuscript, Benson Deverett for helping with the stimulus presentation script in python, Elizabeth McDevitt for suggestions on the figures, Sara Chuang for helping with stimulus selection, and the members of the Hasson, Pillow, and Norman labs for their comments and support. This work was supported by NIMH R01 MH112357-01 to UH and KAN and NINDS D-SPAN award F99 NS118740-01 to JW.

## Notes

### Competing Interest Statement

The authors have declared no competing interest.

